# Task-induced internal bodily rather than brain states regulate human self-perception

**DOI:** 10.1101/2024.11.22.624055

**Authors:** Musi Xie, Han Bao, Hang Wu, Yihui Zhang, Yueyao Liu, Junrong Han, Xilin Zhang, Pengmin Qin

## Abstract

A fundamental ability for us is to identify and distinguish with others, known as the self, whose neural substrate is hard to identify by external self-related stimuli because of possible interference from internal self-related states of both body and brain. Using the same stimuli, we ruled out this dilemma by manipulating human subjects performing own- and celebrity-face discrimination tasks to induce self-related and non-self-related states, respectively. Results showed that stimulus-driven sensory sensory differences among own-, celebrity-, and stranger-faces were independent of task-induced internal states, whereas their perceptual differences were strongly modulated by these states. Intriguingly, we further found that subjects’ bodily (indexed by heartbeat-evoked potentials) rather than brain (indexed by pre-stimulus α-powers) states not only predicated their self-perception but also moderated the relationship between external stimuli and self-perception. Our results reveal for the first time an adaptive self-perception, shaped by not only external stimuli but also internal states, especially the interoception.

## Introduction

Nervous system is well known in a constant internal state of flux that sets our brain in a specific working mode according to perceptual requirements that are updated dynamically and ultimately contribute to flexible behavioral arises (McCormick et al., 2020; Zhang and Xu, 2022). Indeed, a growing body of evidence suggests that perception is not purely constructed by external sensory inputs, but depends on the top-down regulation from various internal states, repeating them across brain regions and modalities to apply similar modulations to different stimuli and species (Gilbert and Li, 2013; Gilbert and Sigman, 2007; Ress and Heeger, 2003; Zhang and Xu, 2022). The construction of a perception involves making the best sense of stimulus sensory inputs based on a set of hypotheses or constraints derived by prior internal states, including bodily aspects like heartbeats (Al et al., 2020; Park et al., 2014), movement (Ayaz et al., 2013; McGinley et al., 2015b), and physiological needs (Seitz et al., 2009)), as well as brain states such as arousal (Han et al., 2024; McGinley et al., 2015a), emotion (Liu et al., 2023; Phelps et al., 2006), expectation (de Lange et al., 2018; Summerfield and Egner, 2009), and attention (Huang et al., 2023, 2020; Martinez-Trujillo and Gulli, 2018; Zhang et al., 2018, 2020). .

In the last two decades, the neural substrate of self, representing our fundamental ability to identify and distinguish with others, is generally identified from neural activity induced by various self-related external stimuli, such as own name (Bao et al., 2023; Liu et al., 2021; Wu et al., 2023; Zhang et al., 2023) or face (Ma and Han, 2012; Ota and Nakano, 2021; Zhou and Jiang, 2015). Considering the self consists of both bodily and non-bodily environmental information, and the importance of an integration between them for self-perception (Qin et al., 2020, 2016; Zhang et al., 2023) which, therefore, is more likely to be strongly modulated by both the bodily and brain states, resulting in the same external stimulus sometimes induces one’s self-perception, but sometimes not. Indeed, evidence from previous electroencephalographic (EEG), magnetoencephalogram (MEG), and functional magnetic resonance imaging (fMRI) studies have demonstrated that human self-perception is shaped by not only the bodily signals (interoceptive information), such as the heartbeats (Aspell et al., 2013; Park et al., 2016; Sel et al., 2016; Suzuki et al., 2013), but also the self-related brain state, for example, the spontaneous activity level in the default-mode network (DMN) links to the ascription of self-relatedness (Qin et al., 2016; Qin and Northoff, 2011). Notably, this internal state-dependent self-perception is also consistent with a recent three-level model of self-processing (Qin et al., 2020), proposing that self-processing, as a gradient organization, on the one hand, could contain three intimately connected levels with different extension: interoceptive-processing, exteroceptive-processing, and mental-self-processing; on the other hand, integrate body-environment information via propagation from interoceptive-processing to mental-self-processing. However, little is known regarding the relative contributions of these two types (body and brain) of internal states to self-perception, and more importantly, whether they independently or interactively shape our self-perception of the world.

Here, to address these questions and to highlight the role of internal states on self-perception, we used the same near-threshold stimuli (own, celebrity, and stranger faces) to maximally (although not completely) reduce stimulus-driven activity (Kouider and Dehaene, 2007; Mudrik and Deouell, 2022). Human subjects in our study were asked to perform Own-Face Discrimination (OFD) and Celebrity-Face Discrimination (CFD) tasks to induce their self-related and non-self-related internal states, respectively (Fig. 1). Besides, EEG signals were acquired during both tasks, and according to previous studies, subjects’ internal bodily and brain states were indexed by the pre-stimulus heartbeat evoked potentials (Park and Tallon-Baudry, 2014) and spontaneous oscillations (Iemi et al., 2017; Samaha et al., 2020; Walz et al., 2015), respectively. Results showed that stimulus-driven sensory differences among own, celebrity, and stranger faces were independent of task-induced internal states, whereas their perceptual differences were strongly modulated by these task-induced internal states. Importantly, we further found that subjects’ bodily rather than brain states not only predicated their self-perception but also moderated the relationship between external stimuli and self-perception. Our results suggest that self-perception is shaped by not only external stimuli but also internal states, especially the interoception.

**Fig. 1.**
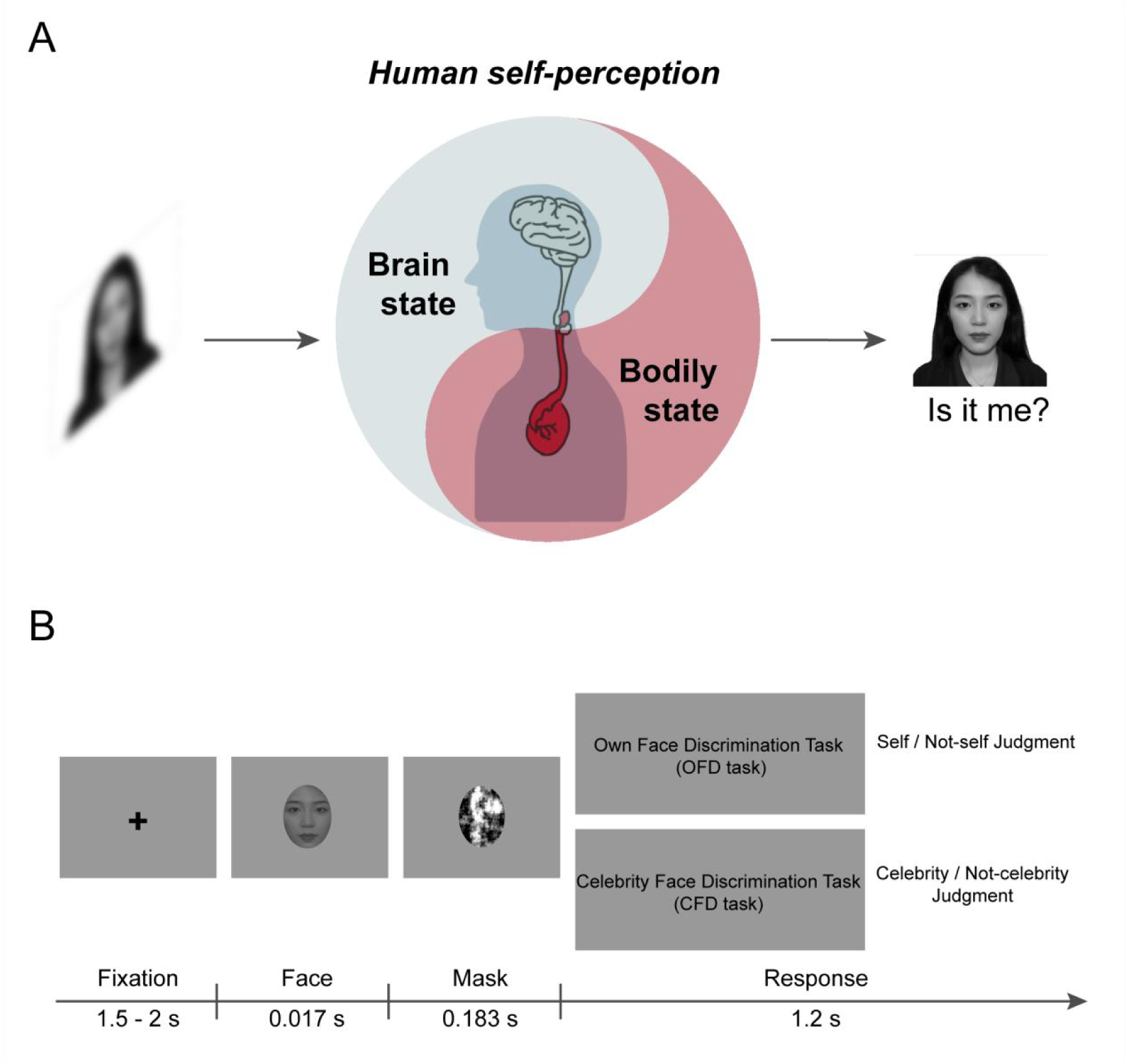
Experimental procedure. (A) Schematic illustration of the human self-perception modulated by internal states (bodily and brain states) (B) Experimental procedure. The subjects were asked to focus on the fixation point at the center of the black screen. A face stimulus, which could be the subject’s own face, a celebrity’s face, or a stranger’s face, was presented for 0.017 s. The presentation of these face stimuli was at equal probability and occurred randomly. This was followed by a mask for 0.183 s and then a fixation cross for 1.2 s. Subjects were instructed to give their answer as quickly as possible with a key press. Each trial was separated by an inter-trial-interval (ITI) marked by a gray screen (0.5 s). The formal experiment comprised two tasks: the Own-Face Discrimination (OFD) and the Celebrity-Face Discrimination (CFD) tasks. In OFD task, subjects were told to judge whether the face was their own or not; in CFD task, subjects were told to judge whether the stimulus was a celebrity or not. The face shown in the figure belongs to one of the authors.

## Results

### Behavioral measurements

Using the same stimuli consisting of one’s own face, a celebrity’s face, and a stranger’s face, human subjects were asked to perform the Own-Face Discrimination (OFD, self- and non-self judgments) and Celebrity-Face Discrimination (CFD, celebrity- and non-celebrity judgments) tasks at near-threshold pedestals, and the order of the two tasks was counterbalanced across subjects. There was no significant difference in either the accuracy (OFD task: 63.4 ± 2.5%, CFD task: 58.6 ± 1.6%, *t_29_* = 1.986, *p* = 0.057) or reaction times (OFD task: 573.08 ± 26.52 ms, CFD task: 564.50 ± 26.07 ms, *t_29_* = 0.566, *p* = 0.576) between OFD and CFD tasks. Moreover, to ascertain that the reported perception was stochastic trial by trial, we analyzed the sequences of judgments by binning the trials into a range of 0 - 10 repetitions (Hesselmann et al., 2008; Rassi et al., 2019). A binomial distribution accounted well for the binned data for both self-judgment in OFD task and celebrity-judgment in CFD task (goodness-of-fit: *R^2^* = 0.974 in OFD task, *R^2^* = 0.916 in CFD task), indicative of no systematic reporting of either judgment.

### Stimulus-driven differences among external stimuli

To reveal the stimulus-driven activity, we computed visual-evoked potential (VEP) evoked by each type of face (i.e., own face, celebrity’s face, and stranger’s face) in both OFD and CFD tasks. The significant differences between own and stranger’s face were evaluated based on cluster-based permutation tests in OFD task based on all the electrode sites (see Methods). The statistical results identified three positive clusters where own face showed higher amplitudes compared to stranger’s face: the first time interval was 140 - 184 ms with the greatest difference at fronto-central electrode sites (cluster sum(*t*) = 2140.2, Monte-Carlo *p* = 0.005, named as VEP1); the second time interval was 292 - 360 ms which was maximal at midline electrode sites (cluster sum(*t*) = 1288.8, Monte-Carlo *p* = 0.013, named as VEP2); the third time interval was 552 - 672 ms (cluster sum(*t*) = 1149.7, Monte-Carlo *p* = 0.016, named as VEP3), maximally observed in the centro-parietal region (Fig. 2A).

**Fig. 2.**
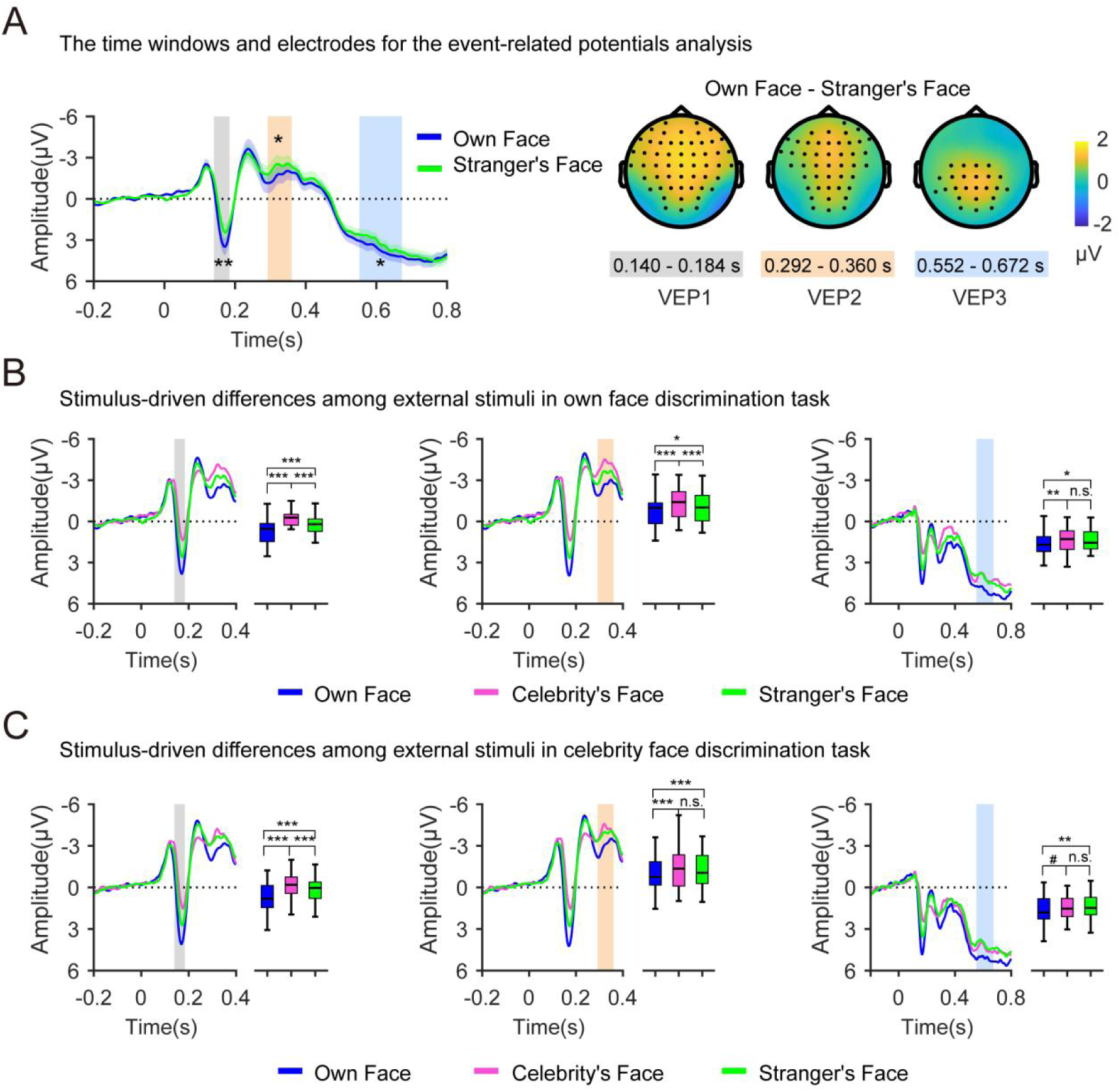
The results of stimulus-driven difference. (A) The left panel shows the time course of the visual-evoked potential (± SEM) for the subject’s own face and stranger’s face in the Own Face Discrimination (OFD) task, averaged over all electrodes. The rectangle area represents the time window in which a significant difference was observed (Monte-Carlo *p* < 0.05, corrected for multiple comparisons in space and time). The right panel showed the topographical map of the difference between own face and stranger’s face conditions in OFD task, grand-averaged in the time window in which a significant difference was observed. Black dots represent the electrodes contributing to the cluster. Averaged across electrodes and time intervals within each of the above clusters, two-factor repeated measures ANOVA, with the task (OFD and CFD) and stimulus (subject’s own face, celebrity’s face, stranger’s face), was performed for each time interval. The time courses and contrasts of OFD task are shown in (B). The time courses and contrasts of Celebrity Face Discrimination (CFD) task are shown in (C). Note: In this figure, the y-axis is reverse ordered. The shaded areas of different colors represent specific time windows utilized for statistical analysis. Each color shade corresponds uniquely to a distinct time window. n.s., not significant, #*p* < 0.1, \**p* < 0.05, \*\**p* < 0.01, \*\*\**p* < 0.001, all *p*-values were corrected for multiple comparisons.

Averaged across electrodes and time intervals within clusters, the mean amplitudes for each type of face were calculated. Then, a two-factor repeated measures ANOVA, with the task (OFD and CFD) and stimulus (own face, celebrity face, and stranger’s face) was performed for each time interval. Results showed that for all three clusters, the main effect of task and the interaction between task and stimulus were not significant (all *p* values are > 0.05); the main effect of stimulus, however, was significant (VEP1: *F_2, 58_* = 47.483, *p* < 0.001, *η_p_²* = 0.621; VEP2: *F_2, 58_* = 25.454, *p* < 0.001, *η_p_²* = 0.467; VEP3: *F_2, 58_* = 9.511, *p* < 0.001, *η_p_²* = 0.247). Post hoc paired t-tests revealed that, for all three clusters, own face exhibited a significant difference in amplitude compared to both celebrity’s and stranger’s face in both OFD and CFD tasks (Fig. 2B and C), except for the comparison between own face and celebrity’s face for VEP3 during the CFD task, which was marginally significant (*p* < 0.10), for detailed statistical results, see Table S1. These results demonstrated that near-threshold own face could evoke distinct activity compared with other near-threshold faces, and these stimulus-driven activities could not be affected by task-induced internal states (self-related or non-self-related). To further verify our conclusions, we conducted a complementary analysis by repeating the above analyses based on the significant difference between own face and stranger’s face in CFD task. These outcomes were consistent with the preceding results. These results support the conclusion that stimulus-driven activities of self could not be regulated by task-induced internal states (self-related or non-self-related), for detailed statistical results, see Fig. S1 and Table S2.

### Perceptual differences among external stimuli

To detect the evoked activity related to subjects’ perception (self judgment, non-self judgment in OFD task; celebrity judgment and non-celebrity judgment in CFD task) of external stimuli, we compared the activities for different judgments. Specifically, two significant perceptual-related clusters were identified in OFD task, the first time-interval was 128 - 188 ms with greatest difference at frontal and central electrode sites (cluster sum(*t*) = 1037.4, Monte-Carlo *p* = 0.001); and the second time interval was 276 - 776 ms, which was maximal from central sites to parietal sites over time (cluster sum(*t*) = 3211.3, Monte-Carlo *p* = 0.018) (Fig. 3A). The time interval within the second cluster was relatively long (500 ms), thus based on the time course of number of electrodes contributing to the cluster over time, we divided the second time interval into two time windows (i.e., 276 - 532 ms and 536 - 776 ms) for the subsequent statistics.

**Fig. 3.**
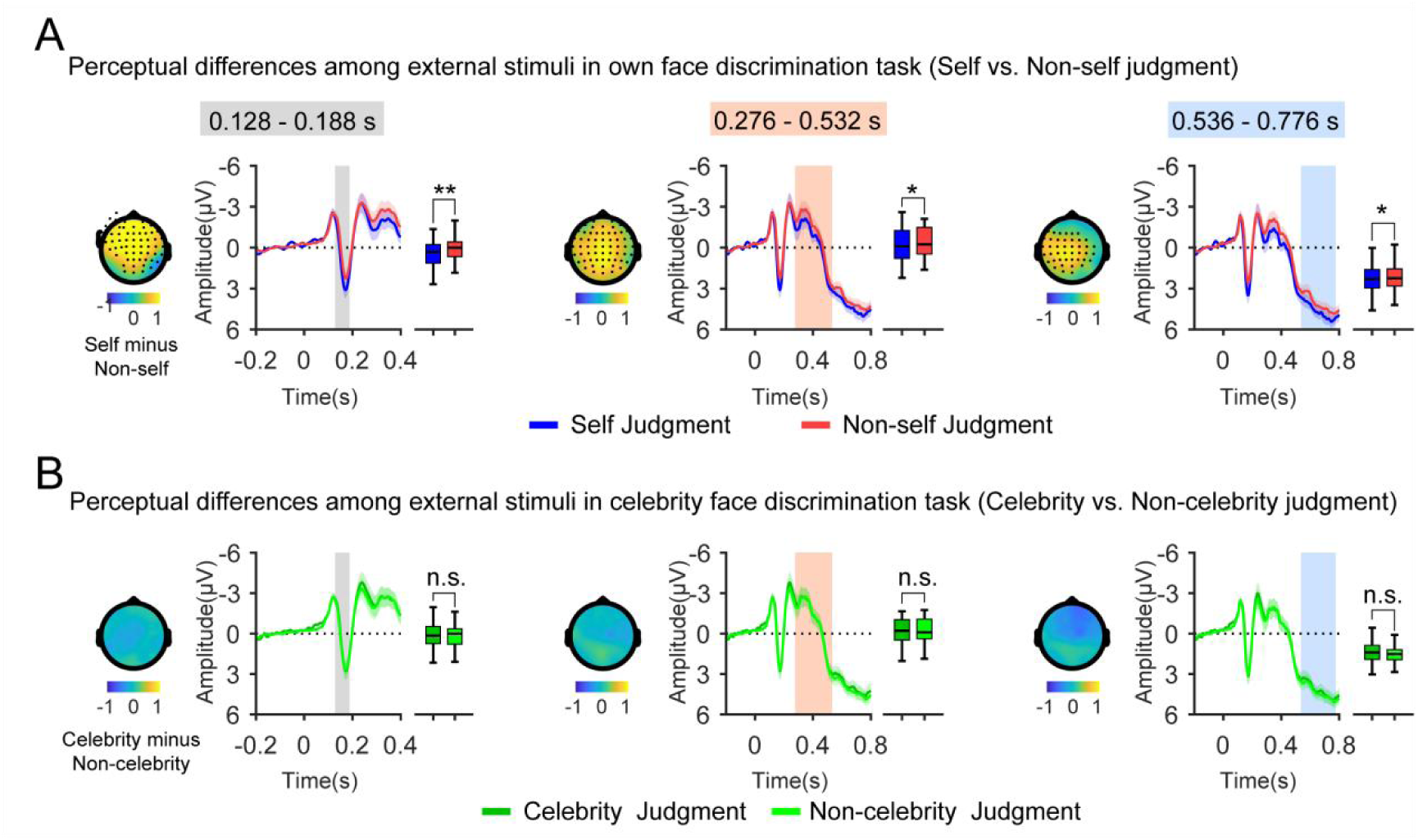
The results of perceptual differences. (A) Perceptual differences among external stimuli in Own Face Discrimination (OFD) task. Time course of the visual-evoked potential (±SEM) for judgment, averaged over the electrodes in each cluster. The rectangular area represents the time window in which a significant difference was observed (Monte-Carlo *p* < 0.05, corrected for multiple comparisons in space and time). Black dots represent the electrodes contributing to the cluster. (B) Perceptual differences among external stimuli in Celebrity Face Discrimination (CFD) task. The mean amplitudes for both celebrity judgment and non-celebrity judgment were obtained by averaging the values across electrodes and the time interval within clusters identified in OFD task, and were compared using a paired t-test. Note: In this figure, the y-axis is reverse ordered. The shaded areas of different colors represent specific time windows utilized for statistical analysis. Each color shade corresponds uniquely to a distinct time window. n.s., not significant, \**p* < 0.05, \*\**p* < 0.01.

In CFD task, we calculated the mean amplitudes for both celebrity judgment and non-celebrity judgment by averaging the values across electrodes and time intervals within clusters found in OFD task. Then, these mean amplitudes between celebrity and non-celebrity judgments were compared using a paired t-test. Crucially, there was no significant difference between celebrity judgment and non-celebrity judgment during each time-window (all *p* values are > 0.05; see Fig. 3B). Additionally, the difference between celebrity judgment and non-celebrity judgment during the whole-time course was also compared by a cluster-based permutation t-test. The non-parametric statistical results showed that there was no significant difference between the two judgments (the smallest Monte-Carlo *p* = 0.065) in CFD task.

### Reduced pre-stimulus alpha power for self-judgment

Since changes in visual-evoked potentials linked to visual perception were observed only in OFD task, we hypothesized that task-induced internal states mediated the effect of self-perception. To investigate whether internal brain states regulate self-perception, we took the pre-stimulus rhythmic nature (i.e., power in delta, theta, alpha and beta frequency band) as an index of the task-induced self-related brain state (Samaha et al., 2020). Compared with non-self judgments, the pre-stimulus activity of self judgments showed a significant reduction in alpha power at central and parietal regions within the time window (from -556 to -16 ms) before stimulus onset (cluster sum(*t*) = -6412.57, Monte-Carlo *p* = 0.018, Fig. 4A). In other frequency bands (delta, theta, and beta), no significant differences were detected (the smallest Monte-Carlo *p* = 0.104).

**Fig. 4.**
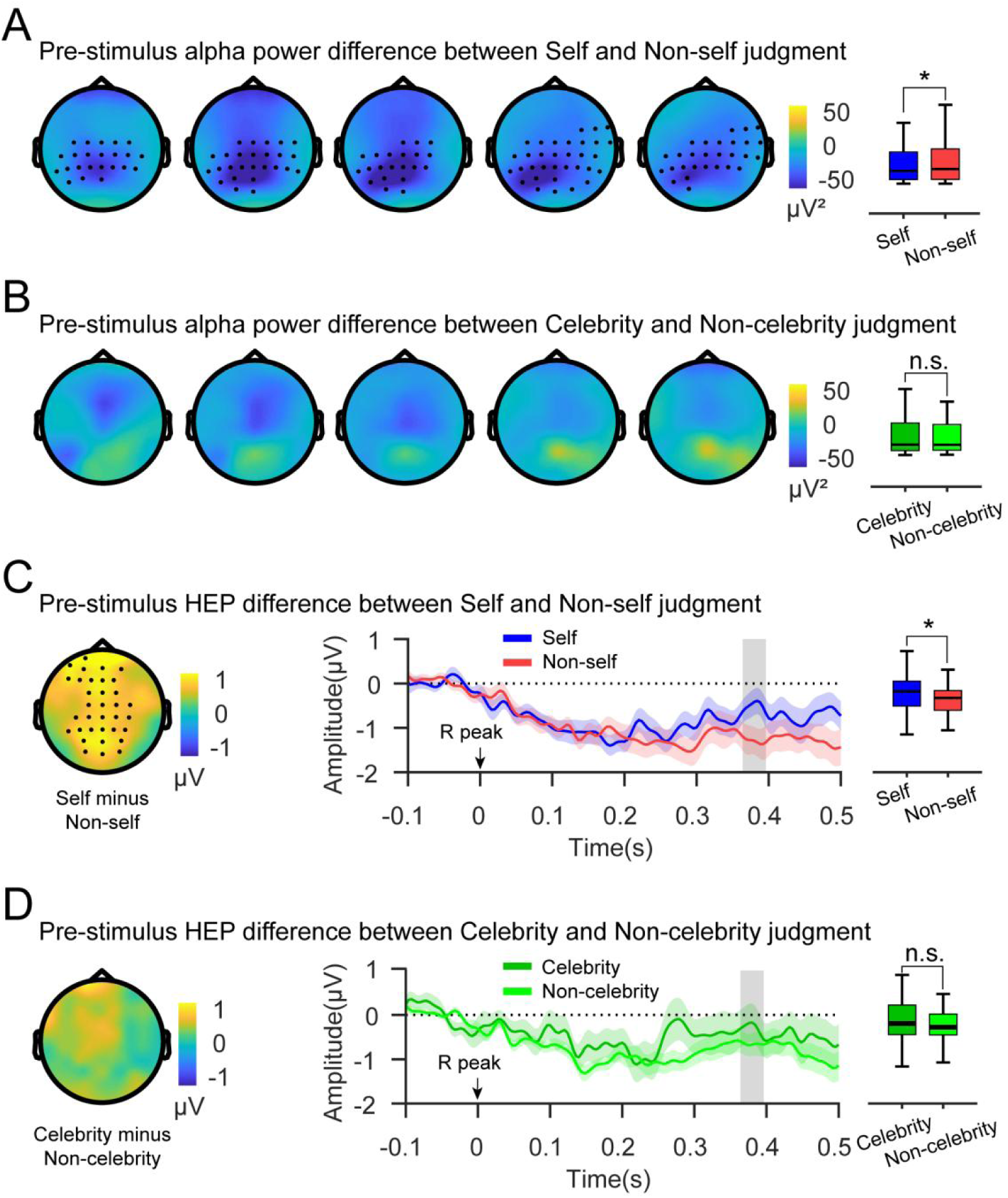
The results of pre-stimulus activity. In the Own Face Discrimination Task (A) and Celebrity Face Discrimination Task (B), the differences in alpha band power before stimulus onset are shown. A significant difference was observed in the Own Face Discrimination (OFD) task from -556 ms to -16 ms before stimulus onset (Monte-Carlo *p* = 0.018), whereas no difference was found in the Celebrity Face Discrimination (CFD) task. The topography maps represent the difference in alpha-band power among judgments (Self - Non-self judgment in the Own Face Discrimination Task and Celebrity - Non-celebrity judgment in the Celebrity Face Discrimination Task). Each map in the series represents the averaged activity over a 108 ms interval within this timeframe, and the black dots indicate the electrodes contributing to the cluster. In the OFD (C) and CFD (D) tasks, the differences in pre-stimulus heartbeat-evoked potential (HEP) are shown. A significant difference was observed in the OFD task between 364 ms and 396 ms (Monte-Carlo *p* = 0.031), whereas no difference was found in CFD task. The topography maps represent the difference in heartbeat-evoked potential among judgments, and the black dots indicate the electrodes contributing to the cluster. Note: n.s., not significant, \**p* < 0.05.

To rule out alternative explanations that the alpha power modulation on judgment in OFD task might purely reflect a general effect associated with Yes/No response, we calculated the mean alpha powers in CFD task by averaging across electrodes and time intervals within the above cluster. The pre-stimulus alpha power between celebrity judgment and non-celebrity judgment showed no significant difference (*t_29_* = -0.525, *p* = 0.603, Fig. 4B). Moreover, in CFD task, the pre-stimulus frequency powers between celebrity and non-celebrity judgments were compared using a cluster-based permutation t-test. The results showed no significant difference in all frequency bands (i.e., delta, theta, alpha and beta, the smallest Monte-Carlo *p* = 0.134). Finally, there was no significant difference in pre-stimulus alpha power between OFD and CFD tasks (*t_29_* = -0.226, *p* = 0.823), indicating that our results cannot be explained by the global changes in brain state between two tasks.

### Increased pre-stimulus HEP amplitude with self-judgment

To investigate whether task-induced internal bodily states regulate self-perception, we took the pre-stimulus HEP as an index to measure the task-induced self-related bodily state (Park and Blanke, 2019; Park and Tallon-Baudry, 2014). In OFD task, a significant difference in pre-stimulus HEP was observed between self judgments and non-self judgments within 364 ms to 396 ms around the midline (cluster sum(*t*) = 443.62, Monte-Carlo *p* = 0.031, Fig. 4C). Similar to the analysis for pre-stimulus power, to rule out the probability that the modulation of pre-stimulus HEP on judgment might reflect a general effect associated with Yes/No response, the mean pre-stimulus HEP amplitudes of celebrity and non-celebrity judgments were calculated and were submitted to a paired t-test. The results showed no significant difference between celebrity and non-celebrity judgments (*t_28_* = 1.393, *p* = 0.175, Fig. 4D). Additionally, in CFD task, the results of the pre-stimulus HEP were also confirmed using a cluster-based permutation t-test, which showed no significant difference (the smallest Monte-Carlo *p* = 0.151). Finally, there was no significant difference in pre-stimulus HEP between OFD and CFD tasks (the smallest Monte-Carlo *p* = 0.103), indicating that our results cannot be explained by the global changes in bodily state between the two tasks.

Correcting the EEG data for the cardiac artifact was performed using an independent component analysis procedure in the data preprocessing step. To eliminate the probability that the observed HEP effect in OFD task was due to cardiac electrical activity (Al et al., 2020; Park et al., 2014), even if the time window analyzed (250 - 400 ms after the R peak) is free from cardiac artifacts (Kern et al., 2013; Schandry et al., 1986). We submitted the ECG signals to a paired t-test. The statistics on the ECG data from 364 to 396 ms after the R-peak did not show any significant difference between self judgment and non-self judgment (*t_28_* = 0.072, *p* = 0.943; see Fig. S2). Furthermore, the results of a paired t-test for each time point showed no significant difference (the smallest *p* = 0.404, with FDR correction across all time points). Taken together, the difference in heartbeat-evoked potential between self judgment and non-self judgment observed on EEG signals cannot be attributed to differences in cardiac electrical activity.

### Logistic regression modeling and permutation importance

After identifying the potential variables — reflecting stimulus-driven activity, bodily state, and brain state — that may contribute to subjects’ self-perception, a multivariable logistic regression model was adopted to identify which variable was most closely related to self-perception. Notably, in this study, subjects were instructed to give their response as quickly as possible; the mean RT in both OFD (573.08 ms) and CFD (564.50 ms) tasks overlapped with the time intervals of VEP3, which ranged from 552 to 672 ms. This suggested that stimulus-driven activity within the VEP3 time interval was unlikely to account for the judgment. Thus, the full model included variables for brain state (indexed by alpha power), bodily state (indexed by HEP), stimulus-driven activity (indexed by VEP1 and VEP2), and their interactions. We first examined whether the predictive performance of the model surpassed chance level (50%). Significance was evaluated using a one sample t-test (one-tailed). The accuracy of OFD task surpassed chance level (58.0 ± 12.7%, *t_28_* = 3.454, *p* < 0.001), while the model showed poor cross-validated performance in CFD task (44.0 ± 15.0%, *t_28_* = -2.138, *p* = 0.979). Second, the performance of the tasks (OFD - CFD) was compared using a paired t-test. Results showed that there was a significant difference between OFD and CFD tasks in terms of accuracy (*t_28_* = 3.507, *p* = 0.002, Fig. 5B).

**Fig. 5.**
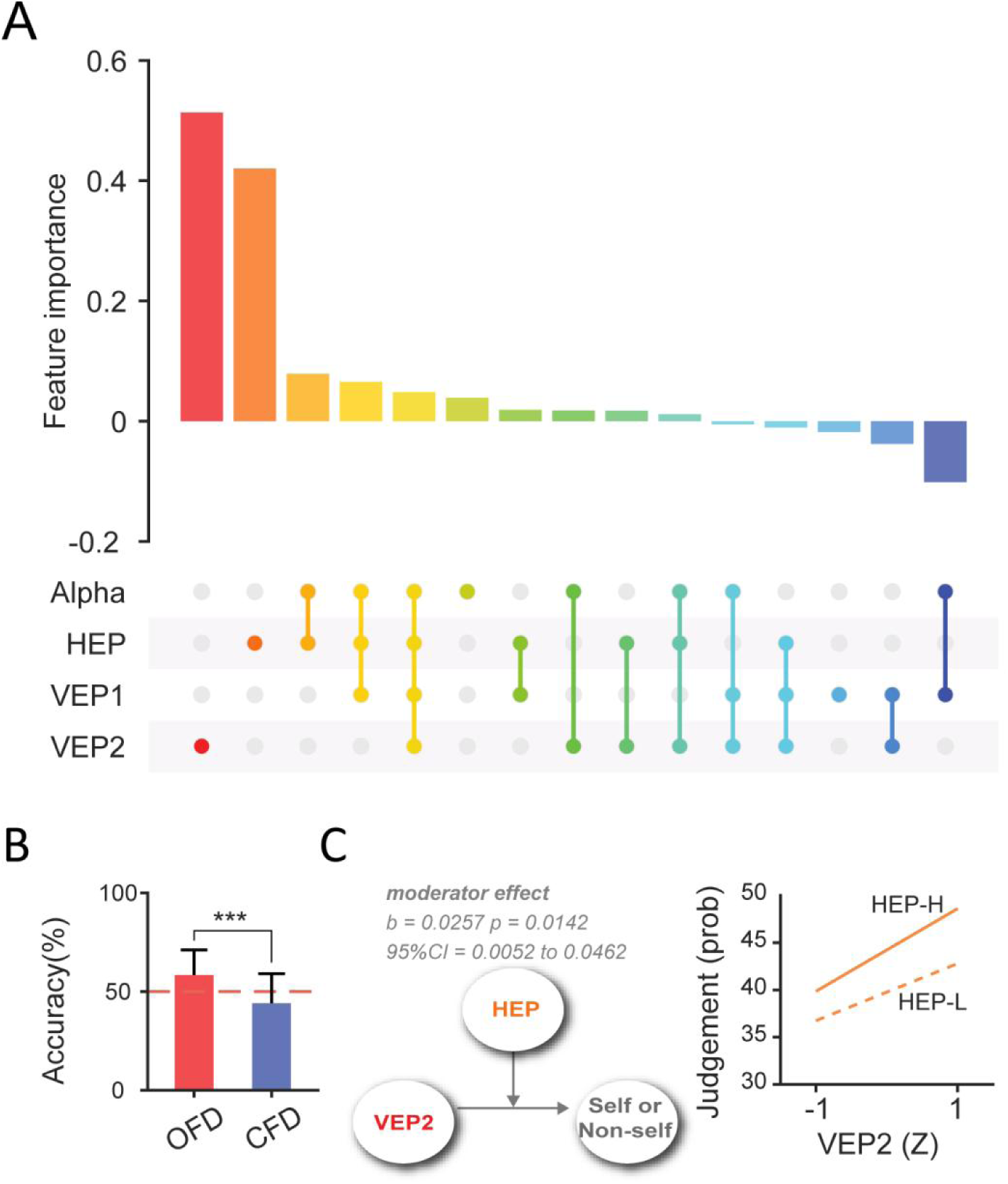
The results of modeling. (A) Feature importance. Each color corresponds to a different variable, providing a visual representation of their relative importance (mean decrease in accuracy). Lines connecting the colored elements represent interactions between the variables. (B) The difference between the Own Face Discrimination (OFD) and the Celebrity Face Discrimination (CFD) tasks in terms of accuracy. (C) Interactive effects of visual evoked potential (VEP2, stimulus-driven activity within the 292-360 ms time interval) and judgment (Self - Non-self judgment in the Own Face Discrimination Task), with heartbeat-evoked potential (HEP) as the moderator. Note: All variables were mean-centered prior to analysis. HEP-H is represented by +1 standard deviation (SD) and HEP-L by -1 SD. Moderator effect significant as 0 not included in the 95% confidence interval. \*\*\**p* < 0.001.

To further evaluate the performance, the F1 score was calculated, which can be defined as the harmonic mean between precision and recall (Manning et al., 2008). The statistical results of the F1 score also revealed a significant difference between OFD and CFD tasks, with F1 score of 72.6 ± 10.4% vs. 49.4 ± 20.3% respectively (*t_28_* = 5.145, *p* < 0.001). Due to the performance of CFD task being at chance level, we did not further explore variable importance analyses. More importantly, the relative importance of the predictor variables in OFD task was calculated by mean decrease in accuracy in the final logistic model, trained on the training set, ranking from large to small. In general, the greater the mean decrease in accuracy when the variable was omitted from the model, the more important the variable. The results revealed that VEP2 (score: 0.513) and pre-stimulus HEP (score: 0.420) were the dominant predictors in our model, as shown in Fig. 5A. The mean values of importance scores for other variables are presented in Table S3.

### Bodily states (HEP) as a Moderator of self-perception

The results in feature importance revealed that HEP was a dominant predictor in comparison with another pre-stimulus variable (i.e., alpha oscillations), and it ranked second in all variables, trailing only VEP2. To further disclose how the self-related bodily state modulates the perception, we performed a moderation analysis with VEP2 as the focal predictor, HEP as the moderator, and judgment as the dependent variable. PROCESS Macro for SPSS with the bootstrap method (moderation *Model 1*) was used to estimate the impact of a moderating variable (A.F., 2012). During the moderation analysis, the interaction term (HEP ×VEP2) was examined to determine whether HEP has a moderating effect on the relationship between VEP2 and judgment. The results showed that the interaction term was significant (*b* = 0.0257, *z* = 2.452, *p* = 0.014; see Fig. 5C and Table S4) When HEP was low (−1 SD), the conditional effect of VEP2 on judgment was positive and significant, when HEP was high (+1 SD), the conditional effect was significant and higher (the value of effect increased from 0.126 to 0.178, Table S5). This indicated that HEP moderated the relationship between VEP2 and judgment (i.e., self-perception). As can be seen from Fig. 5C, the increase in VEP2 will increase the probability of self judgments. The increase in the HEP will strengthen the relationship between VEP2 and self judgments.

## Discussion

The current study for the first time discloses that the self-related bodily state (rather than the brain state), as indexed by heartbeat evoked cortical activity here, regulates human self-perception by modulating the brain’s response to external stimuli. The near-threshold own face evokes distinct activity from other faces (celebrity and stranger), independent of the tasks, indicating specific self-related, stimulus-driven activity. Taken together, the current results demonstrate that human self could contain both flexible (adaptive bodily states-driven) and inflexible (stable external self-related stimulus-driven) characteristics.

Our first main finding is that the own face could evoke distinct potentials from other faces (celebrity and stranger) in both tasks. These findings are supported by previous studies showing that a supraliminal own face could evoke distinct potentials within these three time intervals (Cygan et al., 2022; Geng et al., 2012; Keyes et al., 2010; Ninomiya et al., 1998; Rubianes et al., 2021; Tacikowski and Nowicka, 2010; Żochowska et al., 2021), which seem to reflect different stages of self-related processing. More importantly, our result directly supports the idea that the own face can be distinguished from others even when stimulus-driven activity is reduced under subliminal conditions, building on the findings such as priming effect (Ibáñez et al., 2011; Pannese and Hirsch, 2010) and attention bias (Bola et al., 2021; Fu et al., 2021; Wójcik et al., 2019). Therefore, these findings support a hypothesis about self-related processing (Northoff, 2011). In this hypothesis, self-related processing underlies the basic relation between external signals and the organism, independent of the human’s awareness of the external stimulus. In the current study, self-related stimulus-driven activity may reflect inflexible and static self-related processing.

Our second main finding links task-induced internal state to self-perception. During the own face discrimination task, where a self-related internal state was induced, both brain and bodily states induced by the task can influence self-perception, while such an effect is not observed in the celebrity face detection task. Our results point out that self-perception is not purely constructed by external sensory inputs, but depends on the top-down regulation from various internal states, potentially including both bodily and brain states. Furthermore, these results cannot be explained by general global changes in internal states between two tasks, since neither pre-stimulus alpha power nor HEP showed a difference between these two tasks. As such, self-perception may be affected by the variation of the internal states in a dynamic way. Furthermore, recent findings have shown that the internal brain state can modulate perception and behavior (McCormick et al., 2020; Zhang and Xu, 2022). For instance, the brain state associated with physiological (e.g., locomotor behavior) (Ayaz et al., 2013; Dipoppa et al., 2018; Erisken et al., 2014; Lee et al., 2014) and pharmacological factors (e.g., anesthesia) (Haider et al., 2007; Mitra et al., 2018) can influence the perception of an external stimulus (e.g., detection ratio of the near-threshold stimulus) in both animal and human studies (Benwell et al., 2022; Iemi et al., 2017; Lovett-Barron et al., 2017; McGinley et al., 2015b; Rassi et al., 2019). In human studies, the active task-induced brain state can modulate the perception of the simple physical attributes of a sensory stimulus, such as orientation(Samaha et al., 2017). More importantly, the fluctuation of brain states in self-related tasks can affect one’s perception of one’s own name or own face (Qin et al., 2016; Zhang et al., 2023). Our findings extend those results by succeeded in linking the internal state with self-perception.

More importantly, this study extends the current knowledge of self-perception, demonstrating that human subjects’ bodily rather than brain states predict their self-perception through moderating the relationship between external stimuli and self-perception. This conclusion is supported by the results of logistic regression and moderation analysis. It demonstrates that the internal bodily states contribute to the adaptive self-perception in our daily life, reflecting the flexibility of the human self-perception to external stimulus, which allows individuals to continuously adjust their relationship with the surrounding (natural and social) environment according to actual needs. This result is supported by previous studies. The cardiac information was regarded as the core of the self (Engelen et al., 2023; Park and Blanke, 2019; Qin et al., 2020), and can affect the self-perception of the face (Sel et al., 2016), name (Zhang et al., 2023), rubber hand (Suzuki et al., 2013), and even full body illusions (Aspell et al., 2013; Park et al., 2018) under certain conditions. Thus, the previous studies, which exclusively focused on the fixed relationship between self-perception and self-related stimulus-driven activity, have deviated from the essence of self-perception (Sui and Gu, 2017). Our results clarify the roles of cardiac information in self-perception, and provide direct evidence to support the hypothesis that the brain’s integration of signals from the external environment with those from within the body is fundamental to our sense of self (Engelen et al., 2023; Park and Blanke, 2019; Seth, 2013). Beyond the previous studies, the current findings reveal the specific roles of stimulus-driven activity and self-related bodily state in the generation of self-perception. Furthermore, given that the current results demonstrate an interaction between the bodily state and VEP2 (time interval: 292-360 ms) in influencing one’s response (self-perception), it suggests that the human self-perception for a face may begin within the 292 - 360 ms after the stimulus onset. Finally, the current findings promote our understanding of patients with disorders of self, such as major depression or schizophrenia (Ebisch and Aleman, 2016). Different from previous studies, which aimed to identify the altered self-related stimulus-driven activity in these populations, future studies might also need to pay more attention to their altered self-related internal state.

It is worth noting that, in our study, task-induced bodily rather than brain states dominate human self-perception. There are perhaps a number of factors that have contributed to this and should be considered further. First, the bodily state in our study was indexed by HEP, which is a neural marker of cortical processing of visceral processing (Babo-Rebelo et al., 2019; Kern et al., 2013; Park et al., 2018, 2016, 2014), thus, it may largely reflect brain-body interactions rather than a pure bodily state (Yu et al., 2014; Zeng et al., 2022). This interaction could account for the greater importance of HEP compared to alpha oscillation, which is considered to reflect a pure internal brain state. Therefore, further work is needed using other pure bodily rhythms, such as respiratory-related or gastric-related activity to parse the relative contributions of bodily and brain states in human self-perception. Similarly, pre-stimulus alpha oscillation is taken as a measure of the internal brain state in the current study; it is possible that the alpha oscillation used here is not sensitive enough to capture large-scale brain dynamics, leading us to draw conclusions that the brain state does not dominate self-perception when compared with the bodily state. This makes it essential to consider and investigate further by adopting other measures of internal brain states, like oscillatory synchrony between brain regions (Fries, 2005; Singer, 1993). Second, we used the same near-threshold stimuli (own, celebrity, and stranger faces) to maximally (although not completely) reduce stimulus-driven activity. It is possible that a moderate effect of the bodily state can only be observed when the stimulus-driven activity is reduced. However, it is still not clear whether and how the bodily state moderates self-perception under states with strong stimulus-driven activity, which deserves further study. Finally, the internal states in our study were induced by different experimental tasks, which lasted a relatively long-term period. These task-induced internal states are relatively stable, which might make them more conducive to interoceptive information, because the time course of bodily rhythms is relatively slow (e.g., one or two heartbeats per second). On the contrary, the temporal dynamics of brain activity operate on a millisecond scale. Therefore, we could assume that the effect of the brain state on self-perception could be transiently triggered and missed in the current study. To verify the results of our study, future research should consider the use of other paradigms to generate transient internal states, such as the task-switching paradigm.

In conclusion, the current results showed that self-related bodily states (rather than the brain state) modulate stimulus-driven activity, and thereby contribute to the generation of self-perception. The current results also demonstrate the dissociation between self-perception and self-related stimulus-driven activity. These findings not only enrich the theoretical hypothesis of self and self-perception but also shed light on the investigations concerning patients with various disorders of self.

## Materials and methods

### Participants

Thirty subjects (15 males, aged 18-24) participated in this EEG experiment. All were right-handed with either normal or corrected-to-normal vision, and had no record of psychiatric or neurological disorders. Before the experiment, they provided written informed consent. The study protocols were approved by the human subjects review committee at the School of Psychology, South China Normal University.

### Stimuli

Three types of face stimuli were used: one’s own face, celebrity’s face, and stranger’s face. Before the formal experiment, the photographs of subjects’ own faces were taken using Canon EOS 70D camera against a white background. For the celebrity’s face, participants’ familiarity with various celebrities of the same gender was assessed using a computer-based questionnaire. They rated each celebrity on a five-point Likert scale, ranging from 1 (not familiar at all) to 5 (very familiar). A celebrity with high familiarity ratings was included in the celebrity’s face stimulus for that participant. For the stranger’s face, photos were randomly selected from a list of face images of other participants of the same gender, who were not acquainted with the current participants. All stimuli were developed and presented using Psychtoolbox and MATLAB 2019a (Mathworks). All face pictures were cropped into an oval template and resized to a resolution of 400 × 300 pixels. Stimuli were converted to grayscale and matched on luminance and contrast by using the SHINE toolbox to control for low-level image properties (Willenbockel et al., 2010), 90 Mask pictures were generated using the 2D fast Fourier transform, and the 2D inverse fast Fourier transform. These pictures were subsequently cropped into an oval template and adjusted to match the luminance of the face stimuli. The contrast for all masks was linearly increased at 8 levels by adjusting the SD of intensity from 0 (level 1, most visible) to 140 (level 8, least visible).

### Discrimination Task

Before the experiment, subjects performed a 3-forced choice discrimination task, a mask group with a certain contrast level was adopted to match individual degradation thresholds for face discrimination (Summerfield et al., 2006) (see SI for more details). In the formal experiment, subjects viewed the same face images presented in the same fashion as during thresholding. The formal experiment comprised two tasks: the Own-Face Discrimination (OFD) and Celebrity-Face Discrimination (CFD) tasks, with task order being counterbalanced across participants. In each task, subjects viewed 540 of the same own face, celebrity’s face, and stranger’s face images, presented in the same fashion as during thresholding. Stimuli were backward masked based on the pre-experimentally determined threshold. Images were presented in 6 blocks of 90 trials (30 own face, 30 celebrity’s face, 30 stranger’s face). Each trial began with a central fixation cross presented from 1500 to 2000 ms. Then, a face display was briefly presented (17 ms), followed by a 183 ms mask image and then a fixation cross for 1200 ms. Subjects were instructed to give their answer as quickly as possible with a key press. Each trial was separated by an inter-trial-interval (ITI) marked by a gray screen (500 ms). Responses were made with two fingers (right index finger or right middle finger, counterbalanced across subjects) to a ‘target’ button and a ‘non-target’ button. In OFD task, subjects were told to press the target button if the stimulus was the own face, and the non-target button if it was not; in CFD task, subjects pressed the target button if the stimulus was a celebrity’s face, and the non-target button if it was not.

### EEG acquisition

The EEG signal was recorded at a sampling rate of 1000 Hz using 64 Ag/AgCI electrodes, which were positioned according to the extended international 10–20 system and mounted in an elastic cap, and amplified using an ActiCHamp plus amplifier (Brain Products GmbH, Germany). The signal was high-pass filtered (0.1Hz) online to remove slow drifts. The reference electrode was positioned on FCz, and the ground electrode was placed at the AFz position. The impedance of all electrodes was kept below 5 kΩ throughout the experiment. One additional electrode was positioned to record the ECG.

## Data analysis

### EEG Preprocessing

Data preprocessing and analyses were performed using MATLAB R2019a (The MathWorks, Massachusetts, USA) and in-house scripts based on functions of the open-source toolbox EEGLAB (Delorme and Makeig, 2004). First, the EEG signal was re-referenced to the average of the left and right mastoid electrodes (TP9 and TP10). Then, the signals were filtered using a 50-Hz notch filter, 0.1 Hz high-pass filter, and 40 Hz low-pass filter, and were finally downsampled to 250 Hz for subsequent analyses. Epochs were time locked to the face onset, including data from 1500 ms before stimulus onset to 1400 ms after onset. Independent component analysis (ICA) was used to correct for artifacts in the EEG. Guided by ICLabel’s classification algorithm (Pion-Tonachini et al., 2019), each independent component was then manually checked. ICA components with cardiac field artifact were determined by segmenting ICA components depending on the R-peak of the ECG electrode and visually selecting the components whose time series matched the time course of the R-peak and t-wave of the ECG (Al et al., 2020).For each participant, we excluded, on average, 2.3 ± 0.6 components in OFD task and 2.3 ± 0.4 components in CFD task (e.g., blinks, vertical/horizontal eye movements, and heartbeat-related components). Amplitude rejection (threshold = ± 200 μV) removed trials with residual artifacts after a linear detrend of the signals. 515.7 ± 31.1 trials for OFD task and 515.2 ± 28.8 trials for CFD task remained for further analysis.

### Event-related potentials (ERPs): Stimulus-evoked activity

The EEG data were segmented into 1,000 ms epochs, beginning at 200 ms prior to stimulus onset, and were baseline-corrected using the pre-stimulus interval. Artifact-free epochs were averaged according to trial conditions to calculate an average waveform for each condition for each subject, and grand averages were then obtained by averaging these evoked potentials across subjects.

#### Stimulus-driven activity

To examine the stimulus-evoked activity, the differences between own face and stranger’s face in OFD task were tested using cluster-based permutation implemented in the FieldTrip toolbox, which corrects for multiple comparisons (Maris and Oostenveld, 2007; Robert et al., 2011). Briefly, the procedure includes the following processing steps: A paired t-test was calculated at each electrode and time point separately. Neighboring electrodes in terms of space, that fell below a *p*-value of 0.017 (i.e., 0.05/3, as three pairwise comparisons would be performed in the subsequent analyses) were grouped into clusters. Within each cluster, the sum of t-values was then calculated. A null distribution was generated using permutated data across subjects (1000 permutations). We calculated the maximum cluster-level test statistic, which provided a corrected *p*-value for each cluster. Clusters with a corrected *p*-value < 0.05 (two-tailed) were considered significant. The amplitude of the cluster corresponded to the averaged data across the electrodes and time intervals, showing a significant difference between conditions. Therefore, the electrodes and time intervals within the cluster (*p*-value < 0.05, two-tailed) were selected for subsequent analyses. To further confirm the stimulus-driven effects, a two-factor repeated measures ANOVA, with the task (OFD and CFD) and stimulus (subject’s own face, celebrity’s face, stranger’s face), was performed. Bonferroni correction was used for post-hoc multiple comparisons in ANOVA. Partial eta-squared (*η_n_²*) was computed as a measure of effect size.

#### Judgment (self-perception) related activity

The difference between the two judgments (i.e., self-judgment versus non-self judgment) in OFD task was tested by cluster-based permutation test to determine if the task-induced self-related internal state could regulate perception (corrected *p*-value < 0.05, two-tailed); if there was a significant difference in OFD task, the clusters were applied to the judgment contrast (i.e., celebrity-judgment versus non-celebrity judgment) in CFD task. Briefly, the mean amplitudes across time intervals and electrodes were calculated, then compared via a paired t-test.

### Rhythmic activity

To investigate the relationship between self-perception and internal brain states, the pre-stimulus rhythmic nature of the neural activity was compared. Specifically, EEG signals were subjected to the time-frequency analysis using the Letswave 7 toolbox. Time-frequency transformation was performed in each channel using the complex Morlet wavelet transformation (2 - 40 Hz), and power values were extracted. No baseline correction was applied (Iemi et al., 2017). Statistical analysis was performed based on the band powers calculated in each of the four canonical EEG frequency bands (delta: 2 - 4 Hz, theta: 4 - 7 Hz, alpha: 8 - 12 Hz, beta: 13 - 30 Hz). In OFD task, the difference between the two judgments (i.e., self-judgment versus non-self judgment) for each frequency band during the pre-stimulus period (i.e., ranging from -1500 ms to -4 ms relative to stimulus onset) was tested by a cluster-based permutation test to determine which bands were associated with self-perception under a self-related internal state (corrected *p*-value < 0.05, two-tailed); if there was a significant difference in OFD task, the pre-stimulus time intervals and electrodes within clusters were applied to the judgment contrast in CFD task (i.e., celebrity-judgment versus non-celebrity judgment). Mean power values were calculated, and then compared via a paired t-test.

### HEP: heartbeat-evoked potential

To investigate the relationship between self-perception and internal bodily states, heartbeat-evoked potential (HEP) were used as measure in this study. The cardiac cycles containing a stimulus were selected. R peaks were detected using the peak detection algorithms included in HEPLAB toolbox (Pandelis, 2021), followed by visual correction. In this analysis, one participant was excluded because of a contact problem with the ECG channel during EEG recording. The trials in which the stimulus onset was at least 400 ms after the preceding R-peak were chosen (Al et al., 2020). In this way, the whole epoch was subsequently baseline-corrected by the 100 ms interval prior to R-peak (Gentsch et al., 2019; Marshall et al., 2020, 2018). The pre-stimulus HEP were calculated, which have been reported to occur between 250 and 400 ms after the R-peak (Al et al., 2021, 2020; Haider et al., 2007; Kern et al., 2013; Schandry et al., 1986). Statistical procedures were followed as described previously. A cluster-based permutation test was used to identify clusters where the HEP showed a significant difference between the two judgments in OFD task (*p*-value < 0.05, two-tailed); if there was a significant difference in OFD task, the clusters were applied to the judgment contrast (i.e., celebrity-judgment versus non-celebrity judgment) in CFD task. Briefly, the mean pre-stimulus HEP were calculated across time intervals and electrodes, and then compared via a paired t-test.

### Modeling

After identifying the potential variables that may contribute to self-perception, we used a logistic regression model with the dependent variable being a binary outcome of judgment (i.e., self judgment or non-self judgment in OFD task, celebrity judgment or non-celebrity judgment in CFD task), and the predictor variables included various factors hypothesized to influence these judgments (e.g., stimulus-driven activity before judgment, bodily state, and brain state). Given our hypothesis that the interactions might be important to self-perception, the interactions among predictor variables were also taken into account in this analysis. Logistic regression modeling was conducted using the glm function with the family=binomial argument, sourced from the stats R package (version 4.3.0). Prior to modeling, all data were z scored using scale function in the stats package. In cases where the ratios between the two classes exceeded 1:2, class weights were adjusted inversely proportional to class size to manage the issue of sample imbalance during training (Zadrozny et al., 2003). The equation of the logistic model was as follows:

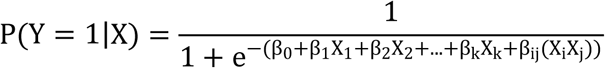

where P(Y = 1 X) is the probability of the dependent variable Y being 1 (i.e., self judgment in OFD task, celebrity judgment in CFD task) given the predictors X. β_0_ is the intercept. β_1_, β_2_,…, β_k_ are the coefficients for the predictor variables X_1_,X_2_,…,X_k_, respectively. β_ij_ represents the coefficient for the interaction between predictor variables X_i_ and X_j_. The logistic function transforms any input into a value between 0 and 1, which can be interpreted as the probability of the instance belonging to the positive class.

To determine the predictive performance of the model regarding judgments and allow for cross-validation of variables’ predictive accuracy, Leave-One-Subject-Out (LOSO) cross-validation was adopted, with accuracy (ACC) serving as the primary evaluation metric. ACC was tested against chance level (50%) using a one-sample t test. If the ACC significantly exceeded chance level (*p*-value < 0.05, one-tailed), the relative importance of various predictor variables within the logistic regression model was quantified using permutation-based feature importance measures. The importance of feature *f* was as follows:

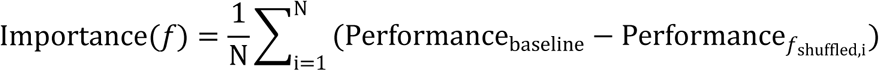

where N is the iteration times; Performance_baseline_ is the performance of the model on the original dataset; Performance*_f_shuffled,i__* is the performance of the model on the dataset where feature *f* is shuffled in the i^th^ iteration. Specifically, 1000 permutations were performed for each variable, systematically reshuffling their values to assess their impact on the model’s performance (i.e., mean decrease in accuracy in test set). The degree to which a model’s predictive performance is degraded by the random permutation of a feature reflects the importance of that feature. Therefore, this approach allowed the identification and ranking of the variables contributing most to the model’s predictive accuracy.

### Moderator analyses

The above modeling analyses showed that HEP was a dominant predictor variable in comparison with another pre-stimulus variable (i.e., alpha oscillations). Among all variables, HEP ranked second, with only VEP2, representing stimulus-driven activity, ranking higher. After identifying the key independent variables based on permutation importance, we proceeded to construct moderator analyses. This analysis aimed to investigate the moderating influence of potential moderator variable on the association between sensation and self-perception. We aimed to understand not only the direct impact of the stimulus-driven activity on the self-perception but also how this impact may be influenced or moderated by other intermediary factors, such as bodily state (i.e., HEP). The dominant pre-stimulus variable detected by importance scores was served as the moderator variable in the model. All data were mean-centred before modeling. Moderation models were tested using PROCESS, a macro developed for SPSS software, version 22 (SPSS Inc.), developed by Andrew Hayes (http://processmacro.org/). In this analysis, the 95% confidence interval (CI) was estimated by bootstrap resampling methods over trials (5000 repeats).

## Author contributions

M.X, H.B, H.W, Y.Z, Y.L, J.H collected data, M.X analyzed data, X.Z and P.Q designed the study and wrote the manuscript, and all authors edited the manuscript.

## Acknowledgments

This research was supported by the National Natural Science Foundation of China (32371098, 32271099 and 31871135); National Outstanding Youth Science Fund Project of National Natural Science Foundation of China (32022032); the Major Program of National Social Science Foundation of China (18ZDA293); and the Key-Area Research and Development Program of Guangdong Province (2019B030335001).

## Data and materials availability

All data are available in the main text or the supplementary materials. Data and analysis files are available at https://osf.io/3zt2k/.

