## Supplementary Materials for "Task-induced internal bodily rather than brain states regulate human self-perception"

Musi Xie *et al.*

 (PQ)

**This Supplementary Materials includes:**

Supplementary Text

Figs. S1 to S2

Tables S1 to S5

### Supplementary Text

### Materials and Methods

#### Discrimination Task

Prior to the discrimination task, subjects performed a 3-forced choice discrimination task on the masking face consisting of 240 trials. Subjects performed this task under identical conditions to the subsequent discrimination task, except that no EEG data were obtained. Images were presented in 8 blocks of 30 trials, and a text prompt reading "1 = one's own face, 2 = celebrity's face, 3 = stranger's face" preceded each block to remind participants of the corresponding response mappings for the thumb, index, and middle finger of their right hand. Images were presented for 17 ms, preceded by a central fixation cross (1500 - 2000 ms), and followed by a randomly selected mask (183 ms). Subjects responded with one of 3 buttons in an uncued fashion following stimulus onset. Masks were initially presented at degradation level 8 (i.e.,which was the least visible with the highest-contrast, intensity SD = 140), in the first block, but a titration procedure was used to keep performance close to 50% accuracy. The accuracy was inspected to determine the contrast level of the mask corresponding to each participant's individual threshold for discriminating the face. The masks with the corresponding contrast level were then used in the discrimination tasks.

**Results**

To further verify our conclusions, we conducted a complementary analysis by repeating the analyses based on the significant difference between own face and stranger's face in Celebrity Face Discrimination Task. These outcomes were consistent with the preceding results. These results support the conclusion that stimulus-driven activities of self could not be regulated by task-induced internal states (self-related or non-self-related). Specifically, the statistical results identified three significant clusters where own face showed a significant difference compared to stranger's face (128 - 192 ms: cluster sum($t$) = 2754.8, Monte-Carlo $p = 0.003$; 272 - 356 ms: cluster sum($t$) = 2593.9, Monte-Carlo $p = 0.003$; 640 - 796 ms: cluster sum($t$) = 2754.8, Monte-Carlo $p = 0.003$). Averaged across electrodes and time intervals within clusters, the mean amplitudes for each type of face were calculated. Then, a two-factor repeated measures ANOVA, with the task (OFD and CFD) and stimulus (own face, celebrity's face, stranger's face), was also performed. Results showed that, for all three clusters, the main effect of task and the interaction between task and stimulus were not significant (all $p$ values are > 0.05); the main effect of stimulus, however, was significant (128 - 192 ms: $F_{2, 58} = 46.720$, $p < 0.001$, $\eta_p^2 = 0.617$; 272 - 356 ms: $F_{2, 58} = 19.826$, $p < 0.001$, $\eta_p^2 = 0.406$; 640 - 796 ms: $F_{2, 58} = 13.098$, $p < 0.001$, $\eta_p^2 = 0.311$). Post hoc paired t-tests revealed that, for three time-intervals, own face exhibited a significant difference in amplitude compared to both celebrity's face and stranger's face, except for the comparison between own face and stranger's face for VEP3 during the OFD task, which was marginally significant ($p < 0.10$), see Table S2.

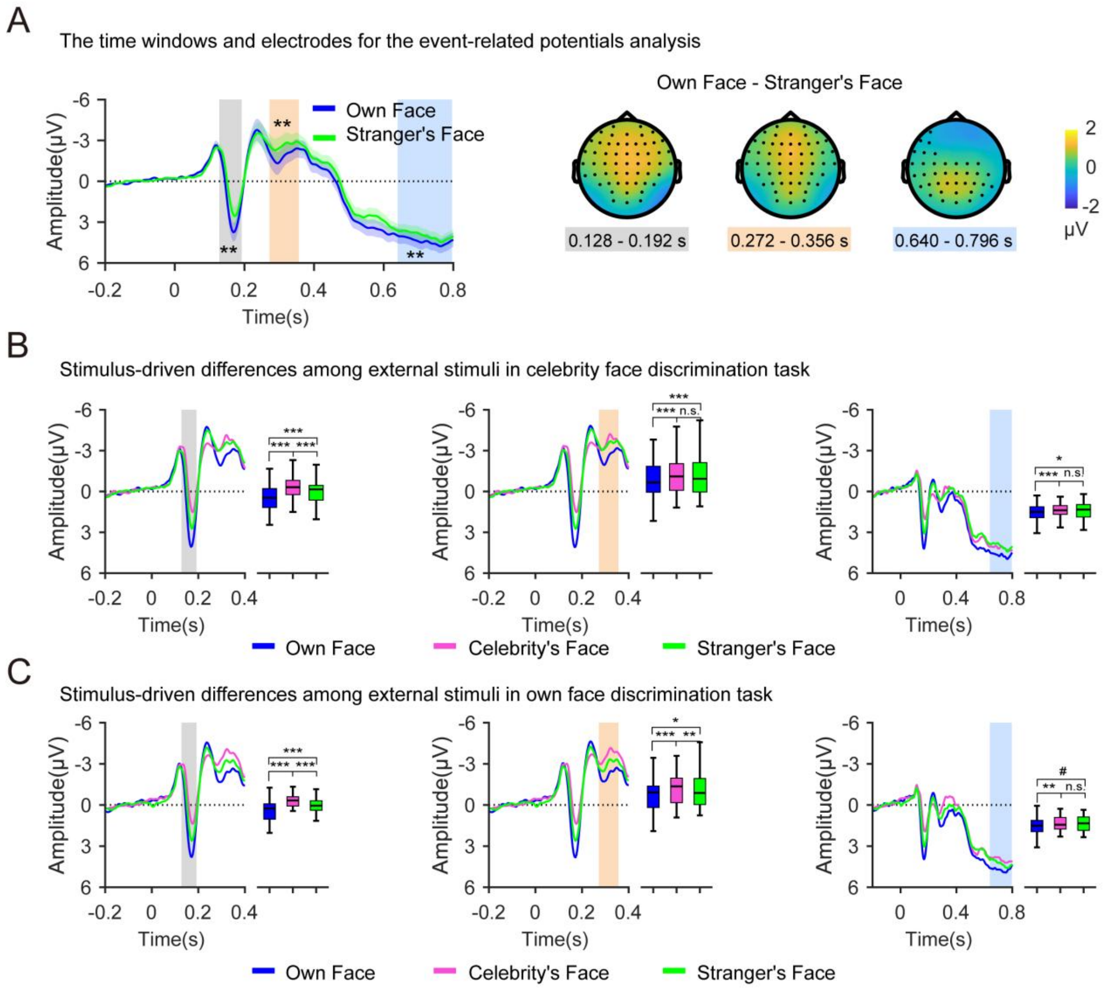


**Fig. S1. The results of stimulus-driven difference** (A) The left panel shows the time course of the visual-evoked potential (± SEM) for the subject's own face and stranger's face in the Celebrity Face Discrimination Task (CFD task), averaged over all electrodes. The rectangle area represents the time window in which a significant difference was observed (Monte-Carlo $p < 0.05$, corrected for multiple comparisons in space and time). The right panel showed the topographical map of the difference between own face and stranger's face conditions in CFD task, grand-averaged in the time window in which a significant difference was observed. Black dots represent the electrodes contributing to the cluster. Averaged across electrodes and time intervals within each of the above clusters, two-factor repeated measures ANOVA, with the task (OFD and CFD) and stimulus (subject's own face, celebrity's face, stranger's face), was performed for each time interval. The time courses and contrasts of CFD task are shown in (B). The time courses and contrasts of Own Face Discrimination Task (OFD task) are shown in (C). Note: In this figure, the y-axis is reverse ordered. The shaded areas of different colors represent specific time windows utilized for statistical analysis. Each color shade corresponds uniquely to a distinct time window. n.s., not significant; #$p < 0.1$; *$p < 0.05$; **$p < 0.01$; ***$p < 0.001$; all $p$-values were corrected for multiple comparisons.

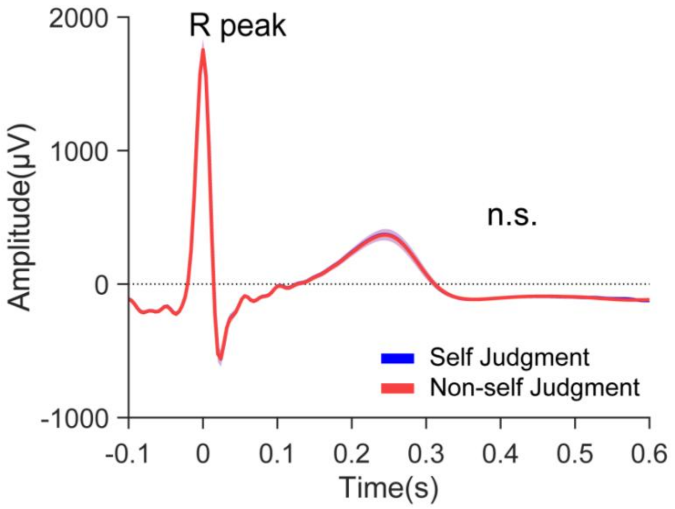


**Fig. S2. R-locked electrocardiogram in Own Face Discrimination Task**. Note: n.s., not significant.

**Table S1.** Results of Post-hoc Pairwise Comparisons for face stimulus based on OFD task

| Time Intervals (ms) | OFD task | | CFD task | |
|---|---|---|---|---|
| | *t* | *p* | *t* | *p* |
| 140 - 184 ms (VEP1) | | | | |
| self vs. celebrity | **8.003** | **< 0.001** | **7.556** | **< 0.001** |
| self vs. stranger | **4.727** | **< 0.001** | **5.234** | **< 0.001** |
| celebrity vs. stranger | **-5.311** | **< 0.001** | **-4.526** | **< 0.001** |
| 292 - 360 ms (VEP2) | | | | |
| self vs. celebrity | **5.698** | **< 0.001** | **5.229** | **< 0.001** |
| self vs. stranger | **3.137** | **0.012** | **4.780** | **< 0.001** |
| celebrity vs. stranger | **-3.968** | **0.001** | -1.239 | 0.676 |
| 552 - 672 ms (VEP3) | | | | |
| self vs. celebrity | **3.282** | **0.008** | 2.453 | 0.061 |
| self vs. stranger | **3.073** | **0.014** | **3.536** | **0.004** |
| celebrity vs. stranger | -0.128 | 1.000 | 1.205 | 0.713 |

Note: OFD task stands for Own Face Discrimination Task, and CFD task stands for Celebrity Face Discrimination Task. Effect with a *p*-value < 0.05 (Bonferroni corrected) was highlighted in bold font.

**Table S2.** Results of Post-hoc Pairwise Comparisons for face stimulus based on CFD task

| Time Intervals (ms) | OFD task | | CFD task | |
|---|---|---|---|---|
| | *t* | *p* | *t* | *p* |
| 128 - 192 ms | | | | |
| self vs. celebrity | **7.789** | **< 0.001** | **7.425** | **< 0.001** |
| self vs. stranger | **4.472** | **< 0.001** | **5.197** | **< 0.001** |
| celebrity vs. stranger | **-5.254** | **< 0.001** | **-4.406** | **< 0.001** |
| 272 - 356 ms | | | | |
| self vs. celebrity | **5.119** | **< 0.001** | **4.388** | **< 0.001** |
| self vs. stranger | **2.771** | **0.029** | **4.511** | **< 0.001** |
| celebrity vs. stranger | **-3.952** | **0.001** | -0.895 | 1.000 |
| 640 - 796 ms | | | | |
| self vs. celebrity | **3.925** | **0.001** | **3.029** | **0.015** |
| self vs. stranger | 2.378 | 0.073 | **4.344** | **< 0.001** |
| celebrity vs. stranger | -1.527 | 0.412 | 1.049 | 0.900 |

Note: OFD task stands for Own Face Discrimination Task, and CFD task stands for Celebrity Face Discrimination Task. Effect with a *p*-value < 0.05 (Bonferroni corrected) was highlighted in bold font.

**Table S3.** Results of permutation importance analysis in OFD task.

| variables | mean | variables | mean |
|---|---|---|---|
| A | 0.039 | V1:H | 0.019 |
| H | 0.420 | V2:H | 0.018 |
| V1 | -0.018 | A:V1:V2 | -0.006 |
| V2 | 0.513 | A:V1:H | 0.066 |
| A:V1 | -0.102 | A:V2:H | 0.011 |
| A:V2 | 0.018 | V1:V2:H | -0.011 |
| A:H | 0.079 | A:V1:V2:H | 0.049 |
| V1:V2 | -0.038 | | |

Note: All variables were z scored prior to analysis. 'A' refers to pre-stimulus alpha power. 'H' refers to pre-stimulus HEP. 'V1' and 'V2' refer to VEP1 and VEP2, respectively, indicating stimulus-driven activity in different time intervals. ':' is used to signify interaction effects.

**Table S4.** Summary of Moderation model.

| | Coefficient | SE | Z | *p* | LLCI | ULCI |
|---|---|---|---|---|---|---|
| constant | -0.3253 | 0.0232 | **-14.0014** | **< 0.001** | -0.3708 | -0.2798 |
| VEP2 | 0.1521 | 0.0242 | **6.2715** | **< 0.001** | 0.1045 | 0.1996 |
| HEP | 0.0915 | 0.0254 | **3.6075** | **< 0.001** | 0.0418 | 0.1411 |
| HEP ×VEP2 | 0.0257 | 0.0105 | **2.4518** | **0.0142** | 0.0052 | 0.0462 |

Note: All variables were mean-centered prior to analysis. LLCI represents the lower level confidence interval, ULCI represents the upper level confidence interval, and SE denotes standard error. Effect with a *p*-value < 0.05 was highlighted in bold font.

**Table S5.** Results of Moderation analysis.

| | Effect | SE | Z | *p* | LLCI | ULCI |
|---|---|---|---|---|---|---|
| -1 SD (HEP) | 0.1264 | 0.0258 | **4.9015** | **< 0.001** | 0.0759 | 0.1769 |
| Mean (HEP) | 0.1521 | 0.0242 | **6.2715** | **< 0.001** | 0.1045 | 0.1996 |
| +1SD (HEP) | 0.1777 | 0.0270 | **6.5773** | **< 0.001** | 0.1248 | 0.2307 |

Note: All variables were mean-centered prior to analysis. LLCI represents the lower level confidence interval, ULCI represents the upper level confidence interval, and SE denotes standard error. Effect with a *p*-value < 0.05 was highlighted in bold font.
